# Multiple endocrine defects in adult-onset Sprouty1/2/4 triple knockout mice

**DOI:** 10.1101/2024.04.18.590049

**Authors:** Gisela Altés, Anna Olomí, Aida Perramon-Güell, Sara Hernández, Anna Casanovas, Aurora Pérez, Juan Miguel Díaz-Tocados, José Manuel Valdivielso, Cristina Megino, Raúl Navaridas, Xavier Matias-Guiu, Ophir D. Klein, Joaquim Egea, Xavi Dolcet, Andrée Yeramian, Mario Encinas

**Affiliations:** Developmental and Oncogenic Signaling Group; Experimental Neuromuscular Pathology Group and; Vascular and renal translational research group; Oncologic Pathology Group, Universitat de Lleida / Institut de Recerca Biomèdica de Lleida, Spain; Department of Pathology, Hospital Universitari De Bellvitge, Barcelona, Spain; Program in Craniofacial Biology and Department of Orofacial Sciences, University of California, San Francisco, San Francisco, CA, and; Department of Pediatrics, Cedars-Sinai Guerin Children’s, Los Angeles, CA, USA

## Abstract

Genes of the Sprouty family (Spry1-4) are feedback inhibitors of receptor tyrosine kinases, especially of Ret and the FGF receptors. As such, they play distinct and overlapping roles in embryo morphogenesis and are considered to be tumor suppressors in adult life. Genetic experiments in mice have defined in great detail the role of these genes during embryonic development, however their function in adult mice is less clearly established. Here we generate adult-onset, whole body Spry1/2/4 triple knockout mice. Tumor incidence in triple mutant mice is comparable to that of wild type littermates of up to one year of age, indicating that Sprouty loss per se is not sufficient to initiate tumorigenesis. On the other hand, triple knockout mice do not gain weight as they age, show less visceral fat, and have lower plasma glucose levels than wild type littermates, despite showing similar food intake and slightly reduced motor function. They also show alopecia, eyelid inflammation, and mild hyperthyroidism. Finally, triple knockout mice present phosphaturia and hypophosphatemia, suggesting exacerbated signaling downstream of FGF23. In conclusion, triple knockout mice develop a series of endocrine abnormalities but do not show increased tumor incidence.

## INTRODUCTION

Sprouty proteins (Spry1-4) are feedback inhibitors of receptor tyrosine kinase (RTK) signaling conserved from flies to humans (Kim and Bar-Sagi, 2004; Cabrita and Christofori, 2008; Edwin et al., 2009; Guy et al., 2009). Homologous recombination experiments in mice firmly establish that during embryonic development the main targets of Sprouty genes are the closely related RTKs, Ret and FGFRs (Mason et al., 2006; Neben et al., 2017). Beyond embryogenesis, aberrant expression of Sprouty family members in different cancer types suggests a tumor suppressive role for this family of genes (Masoumi-Moghaddam et al., 2014; Kawazoe and Taniguchi, 2019).

During development, Sprouty genes act synergistically or independently to ensure proper morphogenesis of the kidney (Basson et al., 2005), inner ear (Shim et al., 2005), enteric nervous system (Taketomi et al., 2005), teeth (Klein et al., 2006), or lung (Taniguchi et al., 2007; Tang et al., 2011), among others. Epistasis experiments show that developmental defects caused by deletion of different family members can be rescued by reducing the genetic dosage of Ret, the FGFRs, or their ligands. For instance, Spry1 knockout mice develop supernumerary kidneys and ureter defects that are rescued upon genetically reducing Ret signaling (Basson et al., 2005; Rozen et al., 2009), whereas Spry2 null mice exhibit diastema teeth that completely disappear upon deletion of Fgfr2 (Klein et al., 2006). Likewise, heterozygous deletion of Spry1 and Spry2 cause aberrant branching of the lung (Tang et al., 2011) and the seminal vesicle (Altés et al., 2022), phenotypes that are completely rescued by concomitant deletion of one allele of FGF10.

Less is known about the role of Sprouty genes in adulthood. Apart from reports demonstrating a role for these genes in regulating homeostasis of different populations of stem cells (Abou-Khalil and Brack, 2010; Urs et al., 2010; Huh et al., 2020), virtually all the literature focuses on their involvement in cancer. Classically, Sprouty genes have been regarded as tumor suppressor genes owing to their inhibitory function (Masoumi-Moghaddam et al., 2014). However it should be noted that in the case of the EGFR, in vitro work supports a role of Spry2 as a positive regulator of the receptor’s function (Haglund et al., 2005; Rubin et al., 2003; Wong et al., 2002) and hence as an oncogene (Kawazoe and Taniguchi, 2019). In any event, most of the above work is based on expression analysis of tumoral vs normal human tissues complemented with functional studies in vitro, with few reports using in vivo techniques and genetically modified mice.

The goal of the present study was to begin to understand the physiological roles of Spry1, Spry2, and Spry4 genes in adult mice through whole-body deletion of these three genes in adult life. We find that tumor incidence is not increased in mutant mice up to one year of age. In contrast, Spry1/2/4 triple knockout mice fail to gain weight over time, show less visceral fat, present mild hyperthyroidism, phosphaturia and hypophosphatemia, and reduced motor activity. In conclusion, we have found multiple endocrine defects but not increased tumor incidence in adult mice lacking all three Spry family members.

## RESULTS

### Efficient whole-body deletion of Spry1, Spry2 and Spry4 in young adult mice

To delete all three Spry alleles in a timely controlled fashion we generated Spry1/2/4 triple floxed mice bearing a Ubiquitin C promoter-driven, tamoxifen-inducible Cre allele [Ndor1^Tg(UBC-cre/ERT2)1Ejb^; (Ruzankina et al., 2007)]. These Cre-expressing animals show widespread recombinase activity upon tamoxifen injection in virtually all organs analyzed (https://www.jax.org/research-and-faculty/resources/cre-repository/characterized-cre-lines-jax-cre-resource). We crossed Spry1^f/f^; Spry2^f/f^; Spry4^f/f^; Ndor1^UBC-cre/ERT2/+^ mice to Spry1^f/f^; Spry2^f/f^; Spry4^f/f^ mice to generate triple floxed mice bearing or not the Cre allele. We then intraperitoneally injected Spry1^f/f^; Spry2^f/f^; Spry4^f/f^; Ndor1^UBC-cre/ERT2/+^ (referred to as “3KO” from this point on) and Spry1^f/f^; Spry2^f/f^; Spry4^f/f^; Ndor1^+/+^ littermates (referred to as “WT”) with 75 mg/kg body weight tamoxifen for five consecutive days at the age of 4-5 weeks. These animals were aged for one year and different measurements taken at defined time points. Animals were then euthanized, and tissues collected (Figure 1a). Efficient deletion of Spry1/2/4 floxed alleles was confirmed in several organs by measuring mRNA levels via RT-qPCR (Figure 1b). There were no differences in the survival rates of both genotypes, with around 80% animals surviving after one year (log-rank χ 0.163 p=0.686) (Figure 1c). Furthermore, when stratified by sex, the Kaplan-Meier analysis revealed no significant differences in all-cause mortality rates between the WT and 3KO groups in males (log-rank χ 0.096 p=0.757) or females (log-rank χ 0.001 p=0.979) (Figure 1d). Gross examination of tissues revealed no signs of tumoral transformation on either genotype. Histological analysis of several tissues including lung, kidney, liver, or thyroid gland showed no evidence of tumoral transformation (Supplemental Figure 1), except for a TTF1-, weakly Calcitonin-positive thyroid tumor in one 3KO mouse, suggestive of medullary thyroid carcinoma (Supplemental Figure 2).

**Figure 1.**
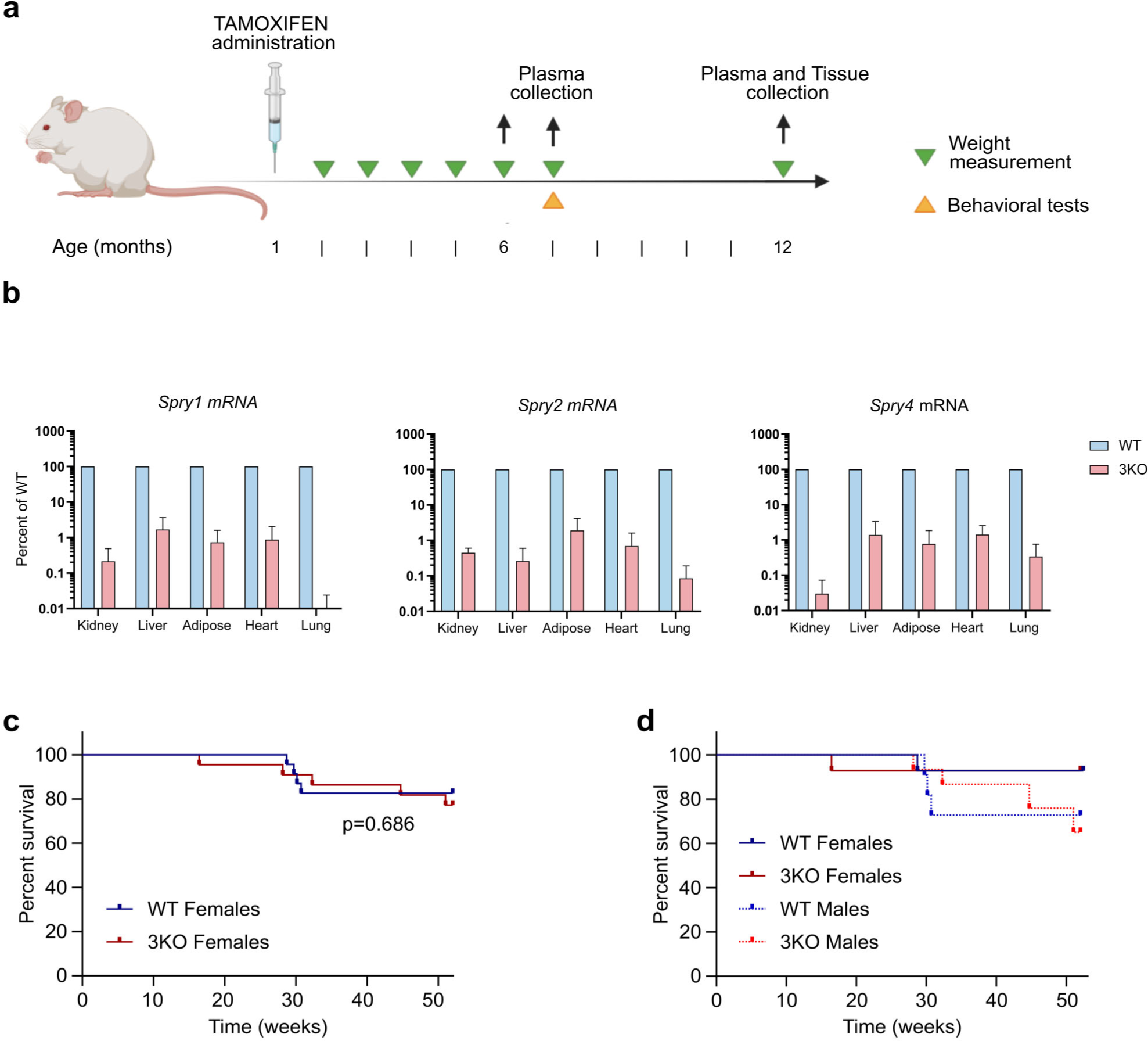
Experimental setup, mRNA expression, and survival analysis. (a) Overview of the schedule of experiments performed with tamoxifen-injected Cre-positive and Cre-negative mice. (b) Analysis of mRNA levels of Spry1/2/4 demonstrates efficient deletion of all three genes in the indicated tissues of 3KO mice. Results are expressed as percent of injected Cre-negative mice (logarithmic scale). Two animals of each genotype were analyzed. (c) KaplanLMeier survival curves reveals no increased mortality of 3KO mice. (d) KaplanLMeier survival curves of the two groups stratified by gender.

### Gross phenotypic changes in 3KO mice

We observed that while WT mice gained weight during the whole period of observation, 3KO animals stabilized their weights roughly two months after tamoxifen injection (i.e. is at around three months of age), with a slight tendency to progressively lose weight after six months of age (Figure 2a, 2b). This behavior was observed both in males (p<0.0001) and females (p<0.0001) and resulted in ∼25-30% weight reduction in both sexes at one year of age (Figure 2b). Interestingly, the amount of visceral fat was drastically reduced in both male and female 3KO mice, being virtually absent in male 3KO mice at one year of age (Figure 2c). Metabolic cage studies revealed no significant differences in food or water intake, as well as urine volume, between WT and 3KO mice, suggesting that alterations in eating habits are unlikely to contribute to the observed differences in weight and fat accumulation between the two groups. Similarly, measurements of physical activity, which we will discuss further later on, do not explain the weight and body fat decrease observed in 3KO mice. In line with the reduced fat levels observed in 3KO mice, basal blood glucose levels were significantly decreased in male 3KO mice at seven months of age (Figure 2g). This finding suggests a potential role of Sprouty genes in regulating glucose metabolism or insulin sensitivity.

**Figure 2.**
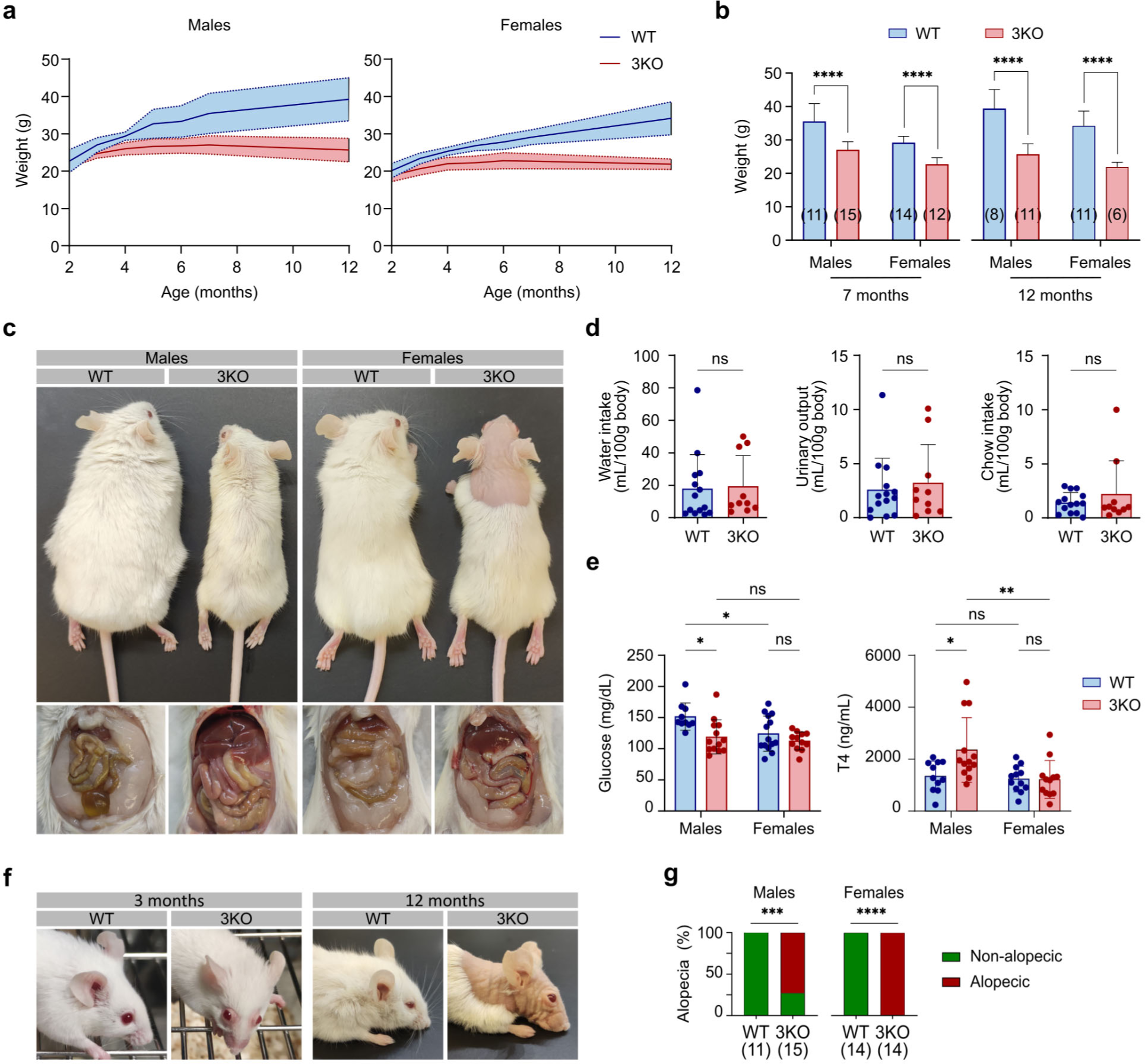
Phenotypic changes of 3KO mice. (a) Unlike WT animals, 3KO mice do not gain weight over time. (b) Weight analysis in WT and 3KO animals by gender at 7 and 12 months of age. (c) Representative pictures of male and female WT and 3KO animals at 12 months of age. In the lower panel, the abdominal cavity is shown, revealing significant differences in visceral fat depots between WT and 3KO animals, both in males and females. (d) Water and food intake, as well as urine volume, are not different between WT and 3KO mice. (e) Plasma glucose and T4 levels of WT and 3KO animals at six months of age. (f) Representative pictures of WT and 3KO mice showing facial abnormalities including alopecia and enlarged eyelids. (g) Incidence of alopecia in 3KO mice by gender at 7 months of age.

By around 3 months of age, 3KO mice presented alopecia around the eyes and nose, including whiskers (Figure 2f and 2g). Hair regrowth was not observed in affected mice housed individually, suggesting that barbering or overgrooming by cage mates was not the cause (data not shown). Alopecia progressed in 3KO mice toward the back of the animals in a continuous, not patchy, pattern. Eyeballs of 3KO mice presented signs of eyelid inflammation (blepharitis) and appeared abnormally protruding (Figure 2f). The eyelid inflammation was characterized by an increase in Meibomian gland size and conjunctival thickness (Supplemental Figures 3 and 4). The weight reduction observed in these animals, coupled with this novel phenotype, rendered the animals readily discernible upon visual inspection. Since we have previously shown that three months old Spry1 knockout mice presented enlarged thyroid glands (Macià et al., 2014), and some of the above phenotypic traits are consistent with hyperthyroidism, we measured plasma levels of T4 of 3KO and wild-type littermates. As shown in figure 2e, T4 levels were significantly higher in male 3KO than in wild type littermates (p=0.0293).

**Figure 3.**
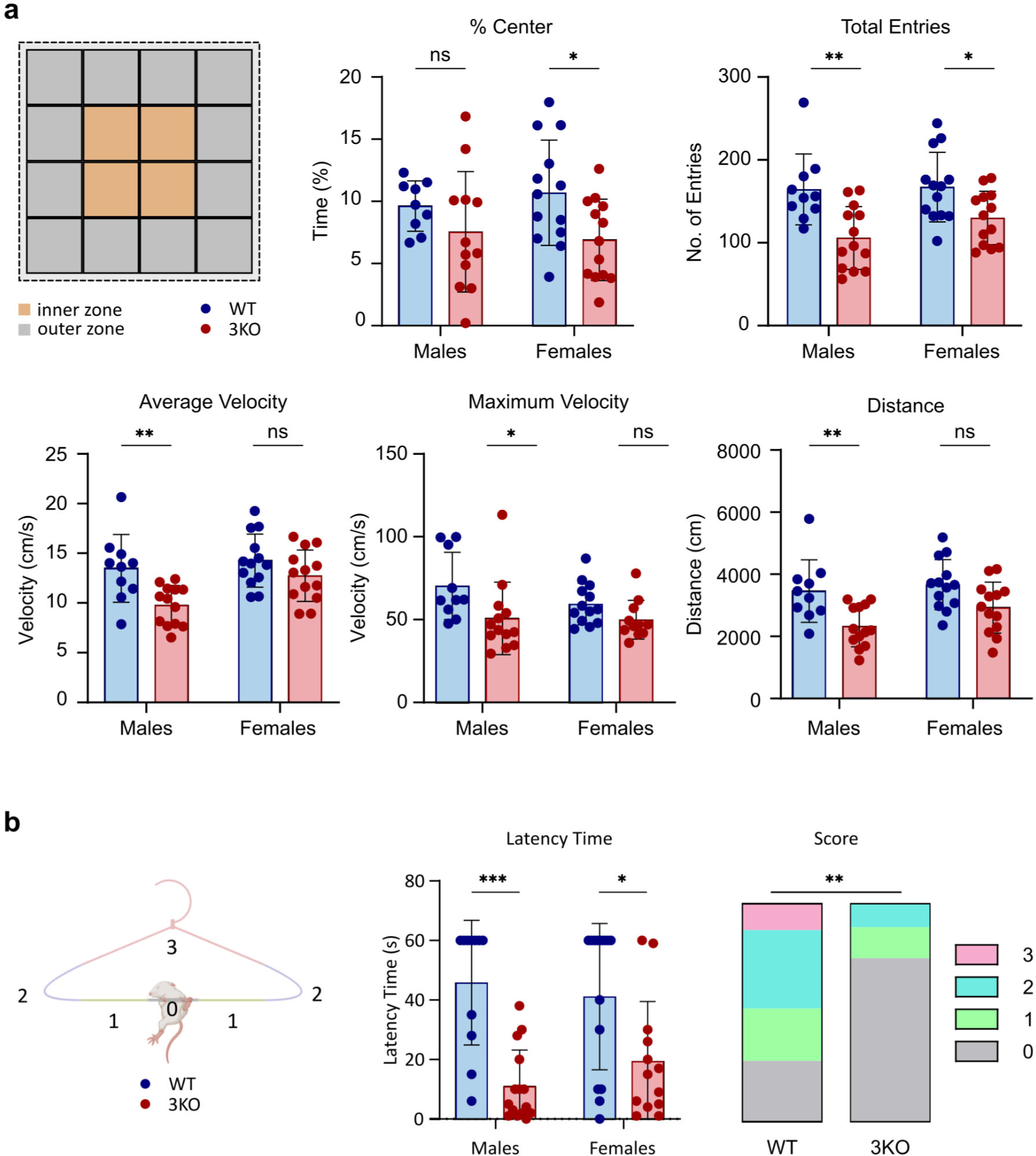
Motor activity deficiencies in 3KO mice. (a) Open-field analysis shows anxiety-like behavior and motor abnormalities in 3KO mice. In the upper left panel a schematic Open-field arena is represented, differentiating central and edge squares. The two following panels show entries to the inner zone (%center), and total entered squares. Bottom three panels show motor performance measured as the average and maximum velocity (cm/s) and distance covered (cm). (b) Coat-hanger test reveals reduced strength and coordination of 3KO mice. In the left panel, the scoring scheme for animal movement is depicted. In the following two panels, the test results are presented, categorized by genotype, with reference to latency time and movement score.

**Figure 4.**
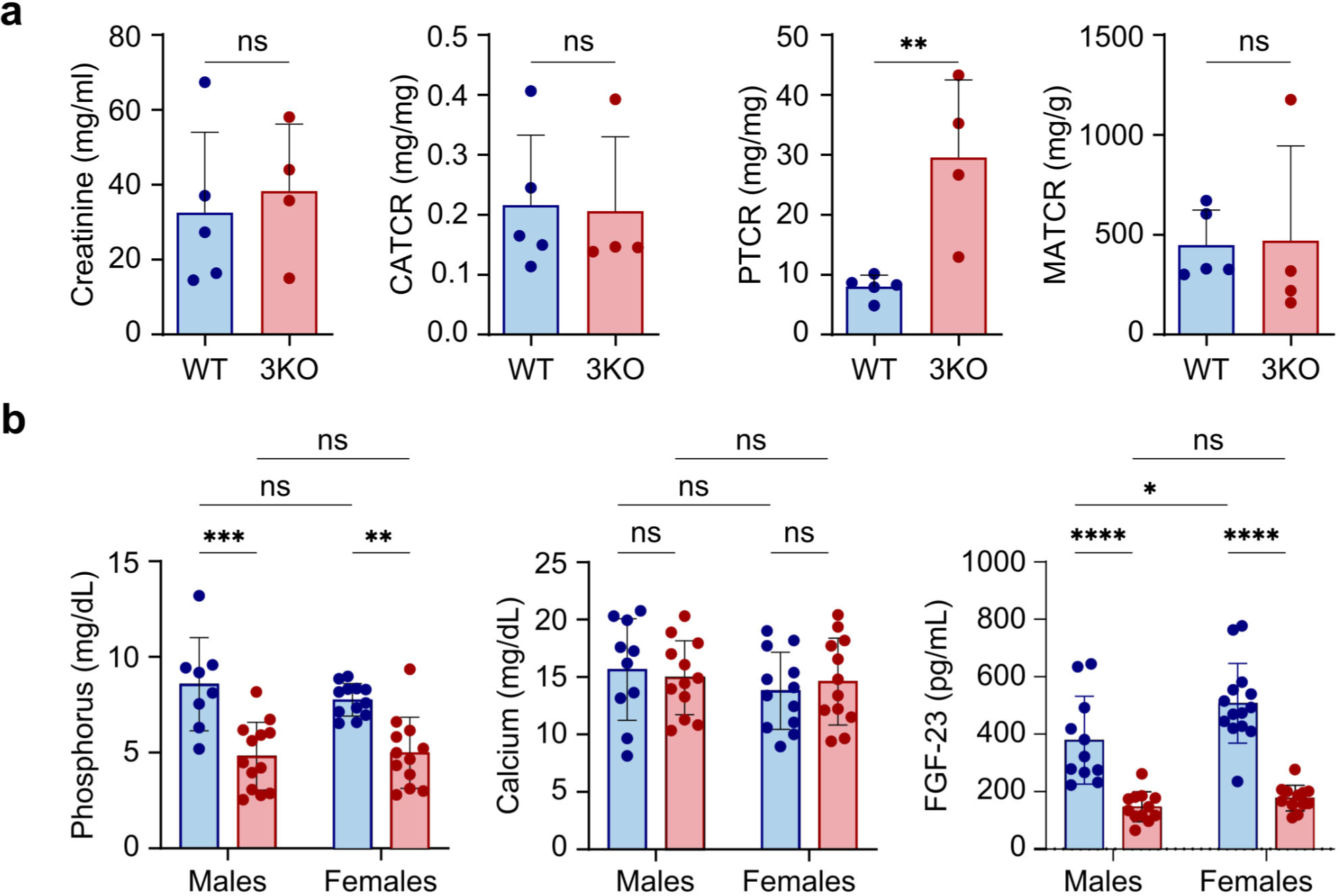
Phosphaturia and hypophosphatemia in 3KO mice. (a) Biochemical analysis of urine metabolites. CATR: Calcium-to-Creatinine Ratio; PTCR: Phosphate-to-Creatinine Ratio; MATCR: Microalbumin-to-Creatinine Ratio. (b) Hypophosphatemia with normal calcium plasma levels of 3KO mice (left and central panels). Right panels show that low phosphorus plasma levels cannot be attributed to abnormally elevated circulating levels of FGF23.

### 3KO animals exhibit increased anxiety and motor activity deficiencies

To begin to undertake a behavioral phenotyping of our mice, we next conducted Open-field tests (Brooks and Dunnett, 2009) on ∼7 months-old wild type and 3KO littermates. When placed in the center of a field, a mouse will typically run to the walled edge and then explore its way around the whole arena while remaining close to the wall. Over time, as the animal habituates to the new environment and its anxiety reduces, the mouse will increasingly venture out towards central parts of the arena before returning to the edges. As shown in Figure3a (top panels), 3KO mice entered fewer times to and stayed shorter times in the inner zone than wild type littermates, suggesting an anxiety-like behavior. Moreover, ambulation measurements showed reduced average and maximum velocity, as well as shorter distances covered by mutant male mice, indicating a reduced motor activity in these mice (Figure 3a, bottom panels). To further evaluate strength and coordination, we also performed the coat hanger test. This test measures the time that mice spend hanging from a wire (usually a coat hanger) before falling (latency time), as well as the regions to which mice climb from the starting point (Figure 3b, left panel). At around 7 months of age, 3KO mice fell more quickly from the coat hanger wire when compared to WT animals (Figure 3b, middle panel). Additionally, 3KO mice demonstrated very few displacements along the bar, indicating weakened grip strength and impaired coordination (Figure 3b, right panel).

### Phosphaturia and hypophosphatemia in 3KO Mice

As previously mentioned, 3KO mice did not show decreased food or water intake, nor they showed decreased urine volume. We next analyzed creatinine, calcium, phosphorus and microalbumin levels in urine, as well as blood urea nitrogen (BUN) from 3KO and wild type mice. Similar microalbumin values in urine (Figure 4a, top panels), as well as comparable BUN levels (Supplemental Figure 5) between genotypes indicated normal renal filtration rates. However, while calcium-to-creatinine ratios were not different, 3KO mice presented significantly higher phosphorus-to-creatinine ratios, indicating specific imbalances in phosphorus management (Figure 4a, top panels). To examine whether phosphaturia inversely correlated with plasma phosphatemia and was not merely reflecting phosphorus overload, we examined both calcium and phosphorus levels in plasma from 3KO and wild type mice. As depicted in Figure 4b, increased phosphaturia correlated with decreased phosphatemia, while calcium levels remained unchanged between genotypes, suggesting abnormal phosphate reabsorption in the kidney of 3KO mice (Figure 4b, bottom left and middle panels). One of the key players in phosphate management is FGF23. This endocrine FGF is released from the bone to the bloodstream, and activates the FGFR-Klotho complex in the kidney tubules to inhibit renal reabsorption of phosphorus. To ascertain whether the phenotype of our mice was caused by high systemic levels of FGF23 leading to impaired reabsorption of phosphate at the kidney, we quantitated plasma FGF23 in both WT and 3KO animals by means of ELISA. As shown in Figure 4b (bottom right panel), we found that FGF23 levels were not only increased, but were drastically reduced in 3KO mice. This indicates that the cause of the observed phenotype was not increased circulating FGF23 in 3KO mice leading to enhanced phosphorus excretion.

## DISCUSSION

In this work we have examined the effects of adult-onset triple deletion of Spry1, Spry2 and Spry4. These 3KO mice present endocrine defects such as hyperthyroidism and low plasma FGF23 levels with hypophosphatemia, and exhibit reduced body weight, visceral fat and motor activity, but do not present an increase in the incidence of tumor formation.

Perhaps the best studied function of Spry proteins apart from embryonic development is its role in tumorigenesis. With the possible exception of Spry2 in colorectal cancer, Sprouty genes are considered to function as tumor suppressors. Downregulation of Spry family members expression has been reported in a long list of human cancers including liver, lung, ovary, or prostate, supporting a role as tumor suppressors (Kawazoe and Taniguchi, 2019; Masoumi-Moghaddam et al., 2014). We have not observed an increase in tumor incidence in 3KO mice up to one year of age. One possible explanation would be that animals analyzed were too young to show differences and indeed no macroscopically visible tumors were found on either genotype besides a putative medullary thyroid carcinoma found in 3KO mice. However, genetic studies in mice show that Spry ablation leads to development of prostate (Schutzman and Martin, 2012; Patel et al., 2013), lung (Shaw et al., 2007), adrenal (Vaquero et al., 2016), and thyroid (Macià et al., 2014) cancers only if a concomitant genetic lesion such as Pten deletion exist. We therefore propose that Spry deletion per se is not sufficient to initiate tumorigenesis but accelerates tumor growth in the presence of driver mutations.

We observed that 3KO do no gain weight over time, and present reduced visceral fat especially in males. Altered adipogenesis might explain this phenotype, as a role of Spry genes in adipogenesis has been described, although data are contradictory and may reflect isoform-specific functions. Thus, on the one hand it has been shown that Spry4 promote adipogenic differentiation from mesenchymal stem cells in vivo and in vitro (Li et al., 2022; Tian et al., 2020). On the other hand, targeted deletion of Spry1 in adipocytes using aP2-Cre mice increases body fat while decreasing bone mass, indicating that Spry1 skews mesenchymal stem cell differentiation towards the osteoclastic lineage (Urs et al., 2010). Our data would be in agreement with a net effect of Spry genes as pro-adipogenic during adipocyte differentiation, but many alternative explanations beyond altered adipocyte differentiation can explain this phenotype. For example, hypersensitivity to FGF21, a weight-loss hormone with multiple metabolic actions such as enhancing insulin sensitivity or promoting lipolysis from white adipose tissue (Flippo and Potthoff, 2021; Zangerolamo et al., 2024) could explain decreased body fat and glycemia of 3KO mice. On the other hand, hyperthyroidism decreases metabolic efficiency by uncoupling oxidative phosphorylation and generating futile metabolic cycles, all of which results in increased thermogenesis and weight loss (Yehuda-Shnaidman et al., 2014). Interestingly, protruding eyeballs (exophthalmos) is a typical symptom of Graves’ disease, an autoimmune hyperthyroidism disorder characterized by overactivation of the TSH receptor by agonistic autoantibodies (Eckstein et al., 2020). Therefore hypersensitivity to TSH could potentially explain both weight loss and exophthalmos of 3KO mice. Since TSH acts via a G protein-coupled receptor, we find doubtful a direct action of Sprouty proteins on its activation. However, at least for orbital fibroblasts, it has been shown that TSH effects are mediated by the IGF1 receptor (Smith and Janssen, 2019), a receptor tyrosine kinase more likely to be a target of Spry proteins. Finally, increases in Meibomian gland thickness could be easily explained by overactivation of the FGFR2(Reneker et al., 2017).

Triple KO mice are hypophosphatemic while showing low plasma levels of FGF23. The most straightforward explanation to this phenotype is that Spry genes attenuate FGF23 signaling at the kidney proximal tubule. FGF23 reduces phosphate renal absorption by inhibiting membrane insertion of phosphate transporters NPT2a and NPT2c via FGFR1 activation of the ERK pathway (Andrukhova et al., 2012; Gattineni et al., 2009). Hyperactivation of FGF23 signaling in 3KO proximal tubule cells would therefore reduce phosphate reabsorption leading to hyperphosphatemia and phosphaturia. Downregulation of systemic levels of FGF23 in this setting could represent a feedback response to low blood phosphate characterizing these animals. Furthermore, indirect mechanisms can also account for hypophosphatemia of 3KO mice. Thus, FGF23 is known to reduce synthesis of calcitriol, the biologically active form of vitamin D, in the kidney proximal tubule via FGFR (Shimada et al., 2004). Hyperactivation of this pathway in the absence of Spry would therefore decrease calcitriol synthesis leading to diminished intestinal absorption and renal reabsorption of phosphate. Moreover, outside the proximal tubule excessive signaling by FGF23 could lead to decreased synthesis of parathyroid hormone (PTH), having a dual effect in phosphatemia since PTH inhibits renal reabsorption in the kidney but in turn promotes phosphate release from bones (Bergwitz and Jüppner, 2010). Finally, 3KO mice also showed reduced motor function and alopecia. This phenotype, together with mineral metabolism alterations found in blood and urine, is reminiscent of that of vitamin D receptor knockout mice, although we have not detected significant decreases in calcium plasma levels. One possible explanation for this would be that calcitriol levels are downregulated owing to overactivation of FGF23 signaling downstream of FGFR in the proximal tubule. Reduced calcitriol activity would lead to alopecia and decreased motor function, with calcium levels being normal due to feedback increase in PTH activity. Studies using tissue specific Cre mice are needed to approach these hypotheses explaining the multiple phenotypes of 3KO mice.

## METHODS

### Animals and experimental protocol

All animal use was approved by the Animal Care Committee of the University of Lleida in accordance with the national and regional guidelines. Mice were maintained on a 12 h light/dark cycle, and food and water was provided ad libitum. Spry1 floxed (Spry1^tm1Jdli^), Spry2 floxed (Spry2^tm1Mrt^) and Spry4 floxed (Spry4^tm1.1Mrt^) mice have been previously described (Basson et al., 2005; Shim et al., 2005; Klein et al., 2006). Ubiquitin C-CreER transgenic mice Ndor1^Tg(UBC−cre/ERT2)1Ejb^ were from the Jackson Laboratories (Bar Harbor, ME, USA). To induce Cre-mediated Sprouty ablation, five consecutive daily intraperitoneal injections of 75 mg/kg tamoxifen were given at 4-5 weeks of age. Tamoxifen (Sigma-Aldrich) was dissolved in absolute 100% ethanol and then diluted to 10 mg/ml final concentration in corn (Sigma-Aldrich). Throughout the course of the study, total body weight measurements were obtained from month 2 to month 12. At 6 months of age, blood samples were collected, and a subset of animals were transferred to metabolic cages to collect urine samples and evaluate their food and water intake patterns. By 7 months of age, behavioral tests were performed. Mice were euthanized at 12 months of age. Blood was collected by cardiac puncture. The organs of interest were collected. One part of the tissues was fixed in 4% paraformaldehyde and the remaining tissue was snap-frozen in liquid nitrogen and kept at −80L°C for protein and RNA extractions.

### Biochemical analyses

Creatinine, phosphorus, and calcium in urine, as well as glucose, phosphorus and calcium in plasma were determined by colorimetric assays (BioSystems). Abuminuria was measured by immunoturbidimetry at the Clinical Analisis Department of the Arnau de Vilanova University Hospital using a Hitachi Modular Analyzer (Roche Diagnostics). To detect FGF-23 plasma levels Mouse/Rat FGF-23 (Intact) ELISA Kit from Quidel (Cat#60-6800) was used. Thyroxine (T4) Competitive ELISA Kit (Invitrogen Cat #EIAT4C) was used to determine plasma T4.

### Open field

To evaluate the effect of Spry deletion on adult motor skills, mice motor performance was periodically tested. The open field test is a widely used behavioral paradigm for evaluating locomotor activity and anxiety in laboratory rodents (Brooks and Dunnett, 2009). Open-field test was performed by the automated recording of mouse movements using Smart Video Tracking software (v2.5.21, Panlab Harvard Apparatus, Holliston, MA, USA) and different parameters such as time, distance, entries in zones, and maximum and average velocity were measured.

### Coat hanger test

In the coat hanger test, mice were placed in the middle of the horizontal portion of a wire coat hanger placed at a height of 40 cm from a table. Mice were released only after firmly gripping the bar with all four paws. The goal was to determine the time the animal stays on the coat hanger (Latency time) and its movement through it (Score) for 60 seconds. The score awarded based on the farthest area the animal has reached from the starting point.

### Histology

Specimens were fixed in 4% paraformaldehyde overnight at four degrees, dehydrated and included in paraffin. Paraffin blocks were sliced at 10 μm, dried for 1 h at 60°C, and then followed a process of dewaxing and rehydration using a xylene/ethanol gradient. Automated Hematoyilin-Eosin staining was performed using a coverstainer device (DAKO).

### RNA isolation and RT-qPCR

RNA isolation was performed using the NucleoSpin®RNA Kit (Macherey-Nagel, Germany) according to the manufacturers’ instructions. RNA was reverse transcribed using the High-Capacity cDNA Reverse Transcription Kit (ThermoFisher) as per manufacturer’s instructions. Quantitative RT-PCR (RT-qPCR) reactions were performed by means of the SYBR green method, using the 2L×LMaster mix qPCR Low Rox kit (PCR Biosystems). Reverse transcriptase-minus and blank reactions were included in all experiments. Serial dilutions of plasmids containing the whole cDNA of murine Sprouty1, Sprouty2 and Sprouty4 were used to generate a standard curve by plotting the number of plasmid molecules vs Ct. The number of molecules in each sample per µg of reverse-transcribed RNA was calculated by interpolating the respective Ct in the standard curve. Expression of each Cre+ sample was Primers used were as follows: Spry1 Fwd 5’-GCGGAGGCCGAGGATTT-3’; Spry1 Rev 5’-ATCACCACTAGCGAAGTGTGGC-3’; Spry2 Fwd 5’-AGAGGATTCAAGGGAGAGGG-3’; Spry2 Rev 5’-CATCAGGTCTTGGCAGTGTG-3’; Spry4 Fwd 5’-GCAGCGTCCCTGTGAATCC-3’; Spry4 Rev 5’-TCTGGTCAATGGGTAAGATGGT-3’

### Statistical Analysis

All data are expressed as means ± standard deviation of the mean. P values and all graphs were generated using GraphPad Prism 9. Statistical significance was calculated using two distinct statistical methods for single variable comparisons, including Mann-Whitney test and two-tailed unpaired t test, depending on normal (Gaussian) distribution of data, studied using Shapiro-Wilk test. To determine the influence of two different categorical independent variables (p.e. genotype and gender) on one continuous dependent variable, two-way ANOVA was used. To compare frequencies Fisher’s exact test was used. The incidence of all-cause mortality was analyzed using Kaplan–Meier curves and the differences between groups were evaluated using the Log-rank test. The level of statistical significance was set at p<0.05.

## ACKNOWLEDGEMENTS

We are grateful to Dr. Shlomo Melmed (Cedars-Sinai Guerin Children’s, Los Angeles, CA, USA) for many suggestions and critical reading of the manuscript. This work was supported by grant PID2020-114947GB-I00, funded by MICIU/AEI/ 10.13039/501100011033, to ME. GA was supported by a predoctoral fellowship from Universitat de Lleida and “Ajuts al Talent en Investigació Biomèdica” fellowship from IRB Lleida. AO is supported by a predoctoral fellowship from Universitat de Lleida. APG is supported by an INVESTIGO contract funded by European Union (NextGenerationEU/PRTR) We thank Marta Hereu, Jessica Pairado and staff from the animal house for their excellent technical assistance.

## AUTHORS CONTRIBUTIONS

Conceptualization, GA, ME; Methodology, GA, SH, AC, JMV, XMG, ODK, JE, XD, AY, ME; Investigation; GA, AO, APG, SH, AP, JMDT, CM, RN, XMG, AY; Formal analysis; GA; Resources; AC, JMV, XMG, ODK, JE, XD, AY, ME; Writing–original draft, GA, ME; Writing–Review & Editing, GA, SH, JMDT, JMV, ODK, JE, XD, AY, ME; Visualization, GA; Supervision, ME; Funding Acquisition, ME.

## REFERENCES

Abou-Khalil, R., and A.S. Brack. 2010. Muscle stem cells and reversible quiescence: the role of sprouty. Cell Cycle. 9:2575–2580. doi:10.4161/cc.9.13.12149.

Altés, G., M. Vaquero, S. Cuesta, C. Anerillas, A. Macià, C. Espinet, J. Ribera, S. Bellusci, O.D. Klein, A. Yeramian, X. Dolcet, J. Egea, and M. Encinas. 2022. A dominant negative mutation uncovers cooperative control of caudal Wolffian duct development by Sprouty genes. Cell Mol Life Sci. 79:514. doi:10.1007/s00018-022-04546-1.

Andrukhova, O., U. Zeitz, R. Goetz, M. Mohammadi, B. Lanske, and R.G. Erben. 2012. FGF23 acts directly on renal proximal tubules to induce phosphaturia through activation of the ERK1/2–SGK1 signaling pathway. Bone. 51:621–628. doi:10.1016/j.bone.2012.05.015.

Basson, M.A., S. Akbulut, J. Watson-Johnson, R. Simon, T.J. Carroll, R. Shakya, I. Gross, G.R. Martin, T. Lufkin, A.P. McMahon, P.D. Wilson, F.D. Costantini, I.J. Mason, and J.D. Licht. 2005. Sprouty1 is a critical regulator of GDNF/RET-mediated kidney induction. Dev Cell. 8:229–239.

Bergwitz, C., and H. Jüppner. 2010. Regulation of phosphate homeostasis by PTH, vitamin D, and FGF23. Annu Rev Med. 61:91–104. doi:10.1146/annurev.med.051308.111339.

Brooks, S.P., and S.B. Dunnett. 2009. Tests to assess motor phenotype in mice: a user’s guide. Nat Rev Neurosci. 10:519–529. doi:10.1038/nrn2652.

Cabrita, M.A., and G. Christofori. 2008. Sprouty proteins, masterminds of receptor tyrosine kinase signaling. Angiogenesis. 11:53–62.

Eckstein, A., S. Philipp, G. Goertz, J.P. Banga, and U. Berchner-Pfannschmidt. 2020. Lessons from mouse models of Graves’ disease. Endocrine. 68:265–270. doi:10.1007/s12020-020-02311-7.

Edwin, F., K. Anderson, C. Ying, and T.B. Patel. 2009. Intermolecular interactions of Sprouty proteins and their implications in development and disease. Mol Pharmacol. 76:679– 691.

Flippo, K.H., and M.J. Potthoff. 2021. Metabolic Messengers: FGF21. Nat Metab. 3:309–317. doi:10.1038/s42255-021-00354-2.

Gattineni, J., C. Bates, K. Twombley, V. Dwarakanath, M.L. Robinson, R. Goetz, M. Mohammadi, and M. Baum. 2009. FGF23 decreases renal NaPi-2a and NaPi-2c expression and induces hypophosphatemia in vivo predominantly via FGF receptor 1. American Journal of Physiology-Renal Physiology. 297:F282–F291. doi:10.1152/ajprenal.90742.2008.

Guy, G.R., R.A. Jackson, P. Yusoff, and S.Y. Chow. 2009. Sprouty proteins: modified modulators, matchmakers or missing links? J Endocrinol. 203:191–202.

Haglund, K., M.H. Schmidt, E.S. Wong, G.R. Guy, and I. Dikic. 2005. Sprouty2 acts at the Cbl/CIN85 interface to inhibit epidermal growth factor receptor downregulation. EMBO Rep. 6:635–641. doi:10.1038/sj.embor.7400453.

Huh, S.-H., L. Ha, and H.-S. Jang. 2020. Nephron Progenitor Maintenance Is Controlled through Fibroblast Growth Factors and Sprouty1 Interaction. J Am Soc Nephrol. 31:2559–2572. doi:10.1681/ASN.2020040401.

Kawazoe, T., and K. Taniguchi. 2019. The Sprouty/Spred family as tumor suppressors: Coming of age. Cancer Sci. 110:1525–1535. doi:10.1111/cas.13999.

Kim, H.J., and D. Bar-Sagi. 2004. Modulation of signalling by Sprouty: a developing story. Nature Reviews Molecular Cell Biology. 5:441–450. doi:10.1038/nrm1400.

Klein, O.D., G. Minowada, R. Peterkova, A. Kangas, B.D. Yu, H. Lesot, M. Peterka, J. Jernvall, and G.R. Martin. 2006. Sprouty genes control diastema tooth development via bidirectional antagonism of epithelial-mesenchymal FGF signaling. Dev Cell. 11:181–190. doi:S1534-5807(06)00255-3 [pii] 10.1016/j.devcel.2006.05.014.

Li, N., Y. Chen, H. Wang, J. Li, and R.C. Zhao. 2022. SPRY4 promotes adipogenic differentiation of human mesenchymal stem cells through the MEK-ERK1/2 signaling pathway. Adipocyte. 11:588–600. doi:10.1080/21623945.2022.2123097.

Macià, A., M. Vaquero, M. Gou-Fàbregas, E. Castelblanco, J.M. Valdivielso, C. Anerillas, D. Mauricio, X. Matias-Guiu, J. Ribera, and M. Encinas. 2014. Sprouty1 induces a senescence-associated secretory phenotype by regulating NFκB activity: implications for tumorigenesis. Cell death and differentiation. 21:333–43. doi:10.1038/cdd.2013.161.

Mason, J.M., D.J. Morrison, M.A. Basson, and J.D. Licht. 2006. Sprouty proteins: multifaceted negative-feedback regulators of receptor tyrosine kinase signaling. Trends Cell Biol. 16:45–54.

Masoumi-Moghaddam, S., A. Amini, and D.L. Morris. 2014. The developing story of Sprouty and cancer. Cancer metastasis reviews. 33:695–720. doi:10.1007/s10555-014-9497-1.

Neben, C.L., M. Lo, N. Jura, and O.D. Klein. 2017. Feedback regulation of RTK signaling in development. Developmental Biology. doi:10.1016/j.ydbio.2017.10.017.

Patel, R., M. Gao, I. Ahmad, J. Fleming, L.B. Singh, T.S. Rai, A.B. McKie, M. Seywright, R.J. Barnetson, J. Edwards, O.J. Sansom, and H.Y. Leung. 2013. Sprouty2, PTEN, and PP2A interact to regulate prostate cancer progression. J Clin Invest. 123:1157–1175. doi:10.1172/JCI63672.

Reneker, L.W., L. Wang, R.T. Irlmeier, and A.J.W. Huang. 2017. Fibroblast Growth Factor Receptor 2 (FGFR2) Is Required for Meibomian Gland Homeostasis in the Adult Mouse. Invest Ophthalmol Vis Sci. 58:2638–2646. doi:10.1167/iovs.16-21204.

Rozen, E.J., H. Schmidt, X. Dolcet, M.A. Basson, S. Jain, and M. Encinas. 2009. Loss of Sprouty1 rescues renal agenesis caused by ret mutation. Journal of the American Society of Nephrology. 20. doi:10.1681/ASN.2008030267.

Rubin, C., V. Litvak, H. Medvedovsky, Y. Zwang, S. Lev, and Y. Yarden. 2003. Sprouty Fine-Tunes EGF Signaling through Interlinked Positive and Negative Feedback Loops. Current Biology. 13:297–307. doi:10.1016/S0960-9822(03)00053-8.

Ruzankina, Y., C. Pinzon-Guzman, A. Asare, T. Ong, L. Pontano, G. Cotsarelis, V.P. Zediak, M. Velez, A. Bhandoola, and E.J. Brown. 2007. Deletion of the developmentally essential gene ATR in adult mice leads to age-related phenotypes and stem cell loss. Cell Stem Cell. 1:113–126. doi:10.1016/j.stem.2007.03.002.

Schutzman, J.L., and G.R. Martin. 2012. Sprouty genes function in suppression of prostate tumorigenesis. Proc Natl Acad Sci U S A. 109:20023–20028. doi:10.1073/pnas.1217204109.

Shaw, A.T., A. Meissner, J.A. Dowdle, D. Crowley, M. Magendantz, C. Ouyang, T. Parisi, J. Rajagopal, L.J. Blank, R.T. Bronson, J.R. Stone, D.A. Tuveson, R. Jaenisch, and T. Jacks. 2007. Sprouty-2 regulates oncogenic K-ras in lung development and tumorigenesis. Genes Dev. 21:694–707. doi:21/6/694 [pii] 10.1101/gad.1526207.

Shim, K., G. Minowada, D.E. Coling, and G.R. Martin. 2005. Sprouty2, a mouse deafness gene, regulates cell fate decisions in the auditory sensory epithelium by antagonizing FGF signaling. Dev Cell. 8:553–564. doi:S1534-5807(05)00079-1 [pii] 10.1016/j.devcel.2005.02.009.

Shimada, T., M. Kakitani, Y. Yamazaki, H. Hasegawa, Y. Takeuchi, T. Fujita, S. Fukumoto, K. Tomizuka, and T. Yamashita. 2004. Targeted ablation of Fgf23 demonstrates an essential physiological role of FGF23 in phosphate and vitamin D metabolism. J Clin Invest. 113:561–568. doi:10.1172/JCI19081.

Smith, T.J., and J.A.M.J.L. Janssen. 2019. Insulin-like Growth Factor-I Receptor and Thyroid-Associated Ophthalmopathy. Endocrine Reviews. 40:236–267. doi:10.1210/er.2018-00066.

Taketomi, T., D. Yoshiga, K. Taniguchi, T. Kobayashi, A. Nonami, R. Kato, M. Sasaki, A. Sasaki, H. Ishibashi, M. Moriyama, K. Nakamura, J. Nishimura, and A. Yoshimura. 2005. Loss of mammalian Sprouty2 leads to enteric neuronal hyperplasia and esophageal achalasia. Nat Neurosci. 8:855–857.

Tang, N., W.F. Marshall, M. McMahon, R.J. Metzger, and G.R. Martin. 2011. Control of Mitotic Spindle Angle by the RAS-Regulated ERK1/2 Pathway Determines Lung Tube Shape. Science. 333:342–345. doi:10.1126/science.1204831.

Taniguchi, K., T. Ayada, K. Ichiyama, R. Kohno, Y. Yonemitsu, Y. Minami, A. Kikuchi, Y. Maehara, and A. Yoshimura. 2007. Sprouty2 and Sprouty4 are essential for embryonic morphogenesis and regulation of FGF signaling. Biochem Biophys Res Commun. 352:896–902.

Tian, L., H. Xiao, M. Li, X. Wu, Y. Xie, J. Zhou, X. Zhang, and B. Wang. 2020. A novel Sprouty4-ERK1/2-Wnt/β-catenin regulatory loop in marrow stromal progenitor cells controls osteogenic and adipogenic differentiation. Metab. Clin. Exp. 105:154189. doi:10.1016/j.metabol.2020.154189.

Urs, S., D. Venkatesh, Y. Tang, T. Henderson, X. Yang, R.E. Friesel, C.J. Rosen, and L. Liaw. 2010. Sprouty1 is a critical regulatory switch of mesenchymal stem cell lineage allocation. FASEB J. 24:3264–3273. doi:10.1096/fj.10-155127.

Vaquero, M., A. Macià, C. Anerillas, A. Velasco, X. Matias-Guiu, J. Ribera, and M. Encinas. 2016. Sprouty1 haploinsufficiency accelerates pheochromocytoma development in Pten+/- mice. Endocrine-related cancer. doi:10.1530/ERC-15-0585.

Wong, E.S.M., C.W. Fong, J. Lim, P. Yusoff, B.C. Low, W.Y. Langdon, and G.R. Guy. 2002. Sprouty2 attenuates epidermal growth factor receptor ubiquitylation and endocytosis, and consequently enhances Ras/ERK signalling. The EMBO journal. 21:4796–808.

Yehuda-Shnaidman, E., B. Kalderon, and J. Bar-Tana. 2014. Thyroid hormone, thyromimetics, and metabolic efficiency. Endocr Rev. 35:35–58. doi:10.1210/er.2013-1006.

Zangerolamo, L., M. Carvalho, L.A. Velloso, and H.C.L. Barbosa. 2024. Endocrine FGFs and their signaling in the brain: Relevance for energy homeostasis. Eur J Pharmacol. 963:176248. doi:10.1016/j.ejphar.2023.176248.

