## Supplemental Figure 1 for "Multiple endocrine defects in adult-onset Sprouty1/2/4 triple knockout mice"

Supplemental Figure 1. Altés et al.

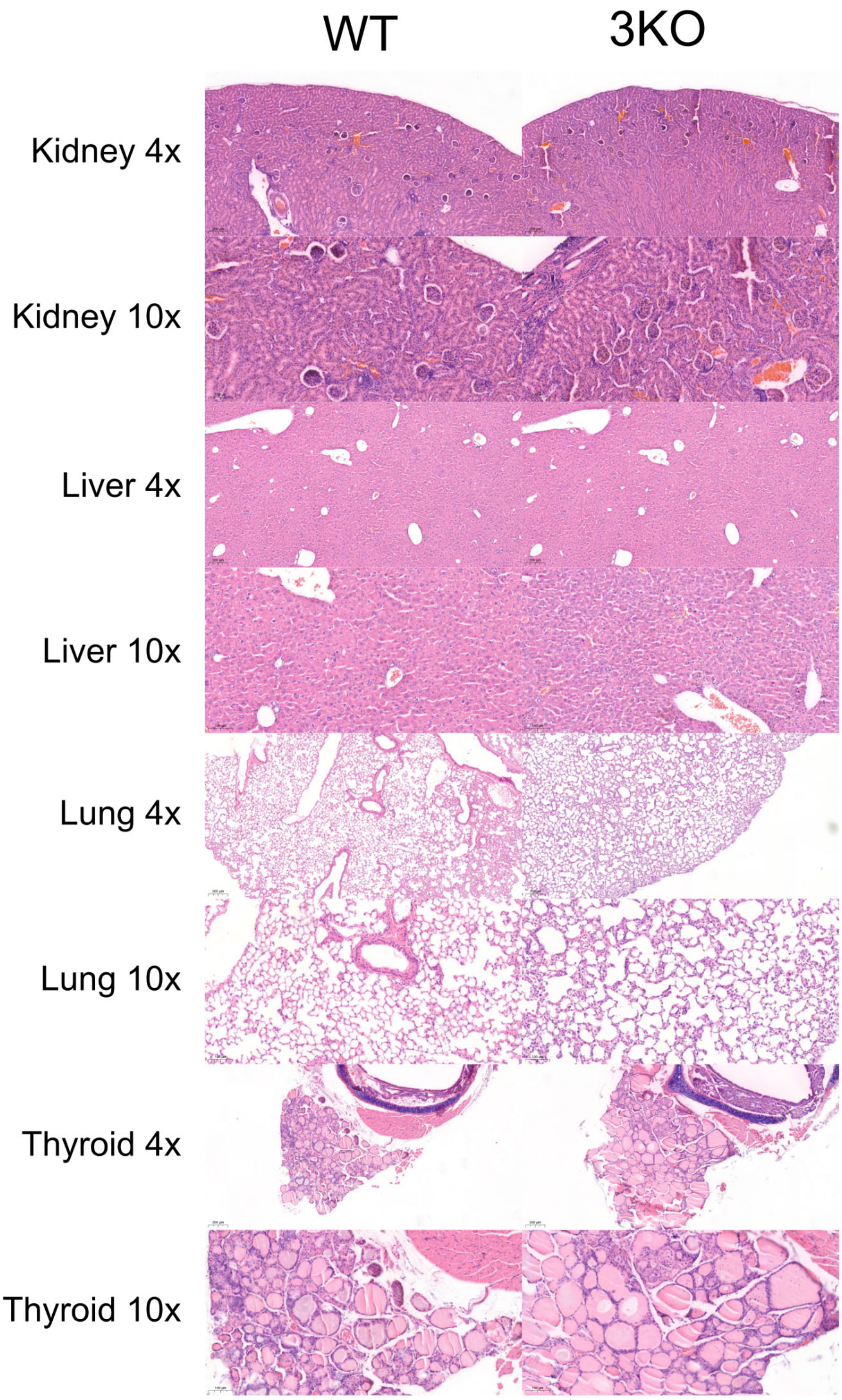


Supplemental Figure 1. Hematoxylin and eosin staining of the indicated tissues from one year old WT and 3KO mice. No neoplastic lesions were observed in tissues from mutant mice.
