## Supplemental Figure 2 for "Multiple endocrine defects in adult-onset Sprouty1/2/4 triple knockout mice"

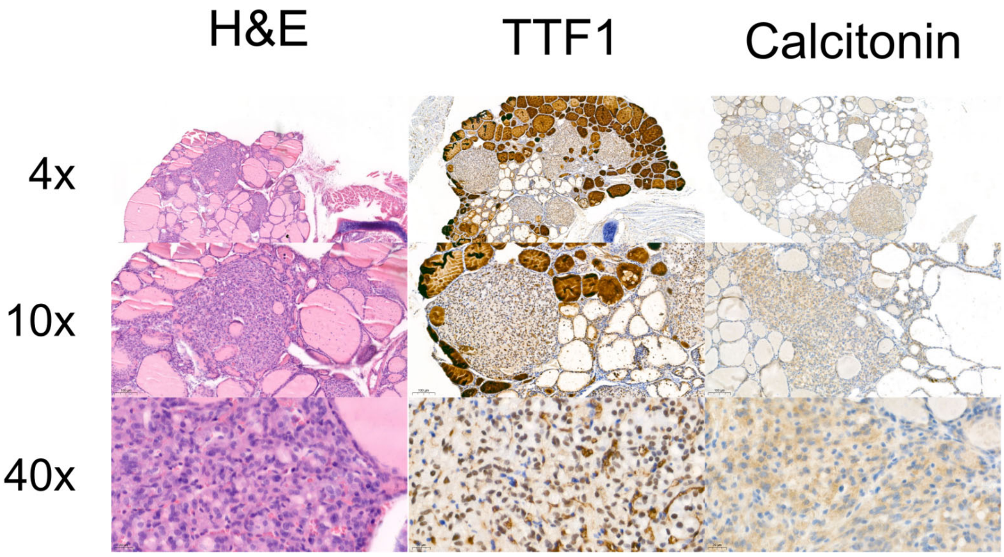


Supplemental Figure 2. Putative medullary thyroid carcinoma found in one 3KO animal at one year of age. Left panels are hematoxylin and eosin staining, middle panel shows positive thyroid-specific transcri ption factor -1 (TTF -1) staining indicating its thyroid origin and right panel shows moderate calcitonin staining, suggestive of medullary thyroid carcinoma. P araffin sections were dewaxed and rehydrated using a xylene/ethanol gradient followed by antigen retrieval (95 °C for 20 min in Tris/EDTA buffer, pH 9) using a PTLink apparatus (DAKO). Staining was performed by an Autostainer device (DAKO)
