## Supplemental Figure 3 for "Multiple endocrine defects in adult-onset Sprouty1/2/4 triple knockout mice"

Supplemental Figure 3. Altés et al.

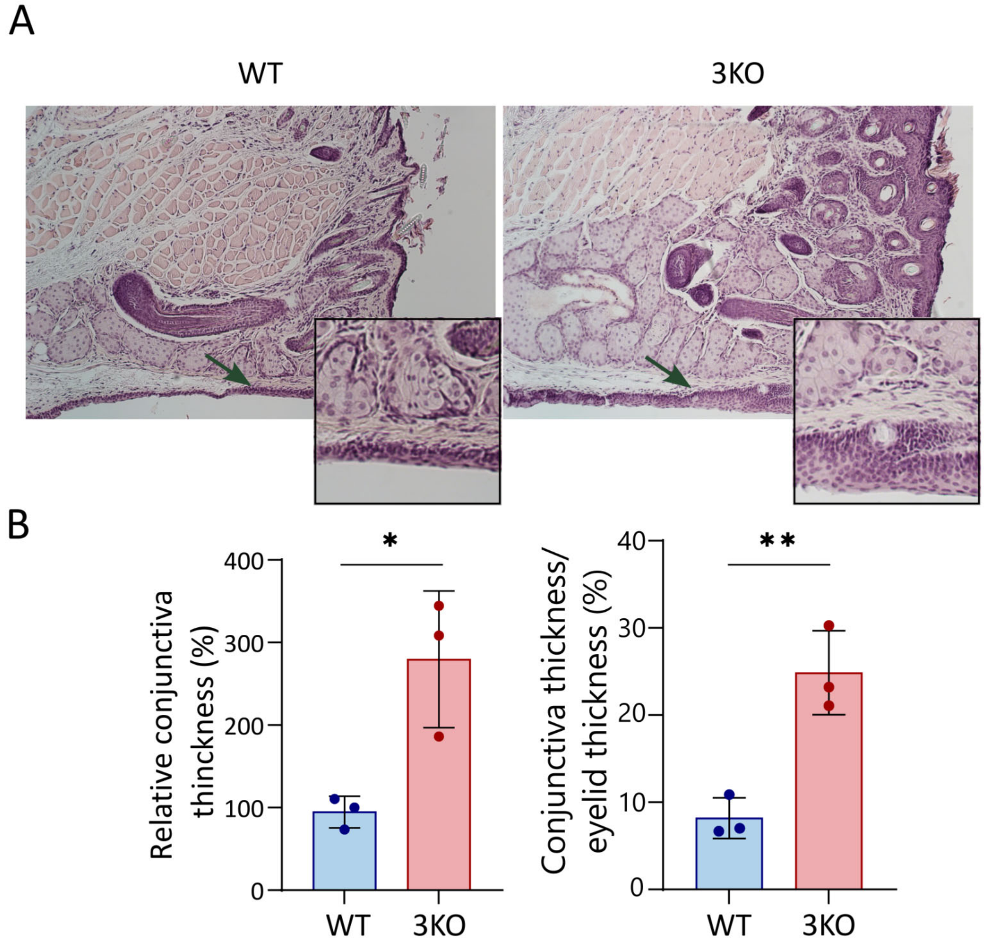


Supplemental Figure 3. Increased conjunctival thickness in one year old 3KO mice. (A) Hematoxylin and eosin staining of eyelids from the indicated genotypes. (B) Quantification of conjunctival thickness. Left panel depicts percent increase relative to WT, right panel shows conjunctival thickness relative to eyelid thickness.
