## Supplemental Figure 4 for "Multiple endocrine defects in adult-onset Sprouty1/2/4 triple knockout mice"

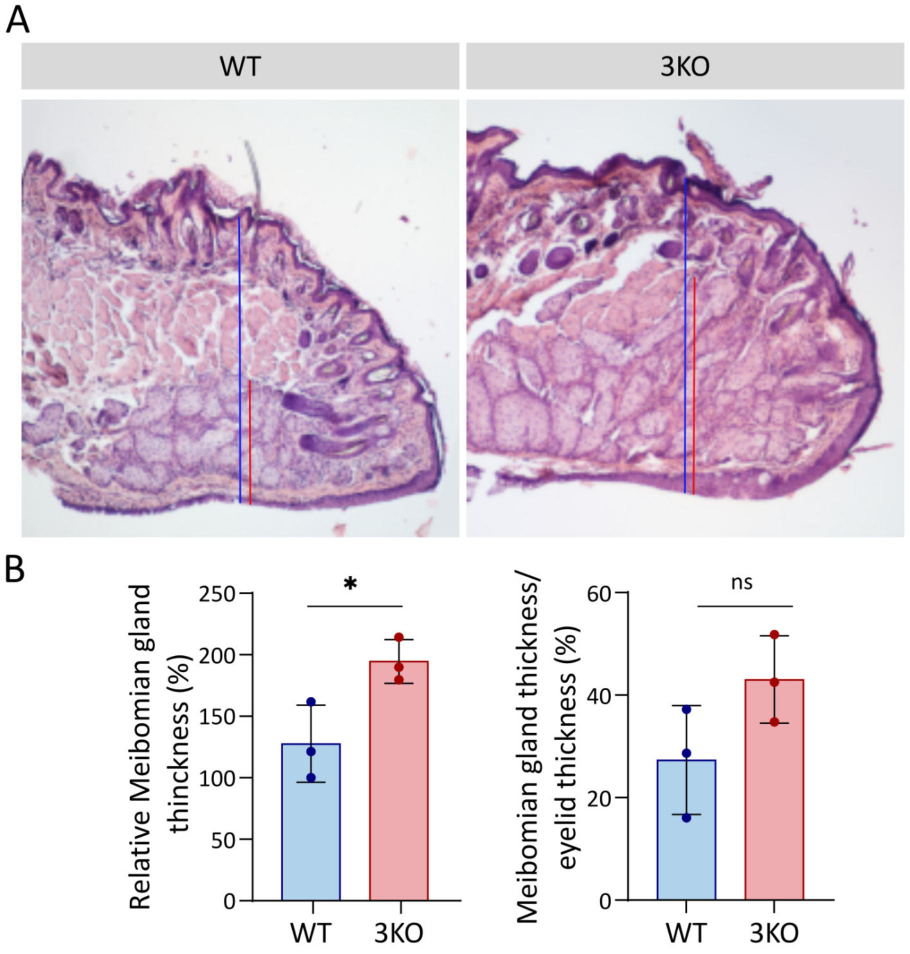


Supplemental Figure 4. Increased Meibomian gland thickness in one year old 3KO mice. (A) Hematoxylin and eosin staining of eyelids from the indicated genotypes. (B) Quantification of Meibomian gland (red line) thickness. Left panel depicts percent increase relative to WT, right panel shows Meibomian gland thickness relative to eyelid thickness (blue line).
