## Supplemental Figure 5 for "Multiple endocrine defects in adult-onset Sprouty1/2/4 triple knockout mice"

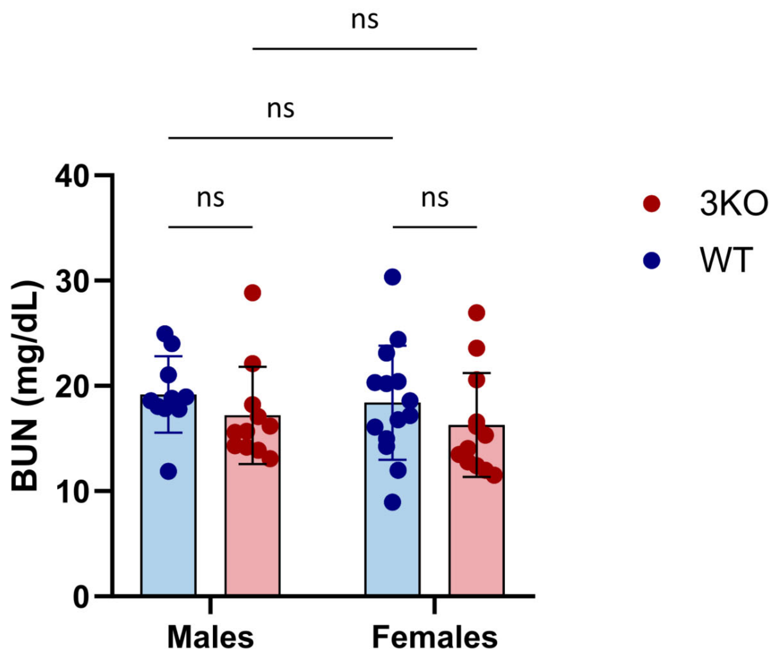


Supplemental Figure 5. Blood urea nitrogen (BUN) levels are unchanged in 3KO mice, indicating normal renal funct on. BUN was determined with a colorimetric assay (Spinreact, Girona, Spain).
